# Redundant neural circuits regulate olfactory masking

**DOI:** 10.1101/2021.04.19.440489

**Authors:** Wenxing Yang, Taihong Wu, Shasha Tu, Myung-Kyu Choi, Fengyun Duan, Yun Zhang

**Affiliations:** Department of Organismic and Evolutionary Biology, Center for Brain Science, Harvard University, Cambridge, MA 02138, USA; Department of Physiology, West China School of Basic Medical Sciences and Forensic Medicine, Sichuan University, Chengdu, Sichuan 610041, China

## Abstract

Olfactory masking is a complex olfactory response found in humans. However, the mechanisms whereby the presence of one odorant masks the sensory and behavioral responses elicited by another odorant are poorly understood. Here, we report that *Caenorhabditis elegans* displays olfactory masking and that the presence of a repulsive odorant, 2-nonanone, that signals threat strongly masks the attraction of other odorants, such as isoamyl alcohol (IAA) or benzaldehyde that signals food. Using a forward genetic screen, we found that several genes, *osm-5, osm-1*, and *dyf-7*, known to regulate the structure and function of sensory neurons played a critical role in olfactory masking. Loss of these genes mildly reduces the response to 2-nonanone and disrupts the masking effect of 2-nonanone. Restoring the function of OSM-5 in either AWB or ASH, two sensory neurons known to mediate 2-nonanone-evoked avoidance, is sufficient to rescue olfactory masking. AWB is activated by the removal of 2-nonanone stimulation or the onset of IAA; however, the mixture of 2-nonanone and IAA stimulates AWB similarly as 2-nonanone alone, masking the cellular effect of IAA. The latency of the AWB response is critical for the masking effect. Thus, our results identify redundant neural circuits that regulate the robust masking effect of a repulsive odorant and uncover the neuronal and cellular basis for this complex olfactory task.

**AUTHOR SUMMARY:** In their natural environment, animals, including humans, encounter complex olfactory stimuli. However, regardless how complex the stimuli are, the behavioral output can only be one. Thus, how the brain processes multiple sensory cues to generate a coherent behavioral output is critical for the survival of the animal. In the present study, we combined molecular cellular genetics, optical physiology and behavioral analysis to study olfactory masking, a common olfactory phenomenon in which the presence of one odorant masks the stimulation of another odorant. Our results show that olfactory masking is regulated by redundant neuronal circuits, a mechanism that likely ensures the robust behavioral response to the sensory cue representing information critical for the survival.

## INTRODUCTION

Odorants represent a wide range of environmental conditions, such as the presence of food, a mate, or a predator. Thus, sensing of odorants regulates behaviors that are critical for survival [1]. The mechanisms through which the nervous system detects an olfactory cue, processes the sensory information, and directs the subsequent behaviors have been characterized in different organisms. Most of these studies use volatile chemicals that are individually presented to signal a simplified environmental condition that is either attractive or repulsive [2–6]. However, attractive and repulsive cues often co-exist under a natural condition where an animal has to make a behavioral decision based on these conflicting messages that respectively signal rewards and threats. Thus, characterizing how the nervous system processes complex odorant information is fundamental for our understanding of how the olfactory system operates.

Previous psychophysics studies on human olfaction show that simultaneous presence of two odorants can evoke a response that is different from the simple addition of the responses to the individual odorants. One of these effects is masking, where the behavioral response to the simultaneous presence of an odorant A and an odorant B is similar to the behavioral response to A or B alone, although A and B each is able to elicit a clear response in isolation [7]. Thus, the presence of one odorant masks the presence of the other. Although the masking effect is widely observed with a profound impact on the perception of a complex olfactory environment, the underlying neuronal mechanism is poorly understood.

While studies on humans and other mammals pioneered the psychophysical analyses of integrated responses to complex olfactory tasks, studies using simple animals, such as *Drosophila melanogaster* and *Caenorhabditis elegans*, utilize complementary approaches to address the underlying molecular and cellular basis [8–12]. The nervous systems of these organisms are highly accessible by genetics, which allows the dissection of the signaling pathways and the neural circuits that regulate odorant-guided behaviors in complex olfactory environments. For example, a fermenting fruit releases multiple odorants, including attractive odorants that signal food and aversive CO_2_, that are detected by fruit flies. A previous study shows that a mushroom body lobe region in the central nervous system of a fruit fly integrates the food odorant - elicited signals to suppress the CO_2_ - evoked output signals. This integration allows the fly to approach food even when the innately repulsive CO_2_ is present [12]. *C. elegans* is strongly attracted by an odorant diacetyl that signals food; however, the attraction can be blocked by a barrier of a hyperosmotic solution. It is shown that tyramine released by an interneuron inhibits an osmosensory neuron that triggers avoidance in response to hyperosmotic stimulation. Suppressing the inhibitory pathway using genetic mutations or food deprivation motivates the worms to cross the hyperosmotic barrier in order to reach the food odorant [11]. Together, these studies demonstrate that the nervous systems of simple organisms use conserved signaling mechanisms to modulate the function of neural circuits to generate a coherent behavioral response when facing sensory cues of conflicting valences.

In this study, we use *C. elegans* to characterize the molecular and neuronal basis of olfactory masking. With a compact and well-defined nervous system [13], *C. elegans* responds to a large number of attractive or repulsive odorants. Several pairs of sensory neurons detect these odorants using G-protein coupled receptors and the downstream cGMP-mediated signaling pathway [2, 14–19]. Among the well-studied olfactory cues, we chose 2-nonanone and isoamyl alcohol (IAA) to make a pair of odorants with conflicting valences. 2-nonanone is a strong repellent that is sensed by the olfactory sensory neurons AWB and ASH. A direct contact with a solution with the high concentration of 2-nonanone kills a worm acutely. Thus, 2-nonanone likely represents a threat to survival [4, 20–22]. In contrast, IAA at low concentrations strongly attracts a worm and is mainly detected by the AWC sensory neurons [2, 23, 24]. In our behavior paradigm, we put 2-nonanone and IAA side-by-side to stimulate the worms with an attractive odorant and a repulsive odorant simultaneously. We find that while IAA alone strongly attracts the worms and 2-nonanone alone strongly repels them, IAA and 2-nonanone together repel the worms as much as 2-nonanoe alone. The attractive effect of IAA is completely masked by the presence of 2-nonanone. By screening for mutants that were generated by random mutagenesis, we identified three genes, *osm-5, osm-1*, and *dyf-7*, which led us to uncover the circuit mechanisms underlying olfactory masking.

## RESULTS

### A new behavior paradigm to characterize olfactory masking

To study the mechanisms underlying olfactory masking, we developed a behavior paradigm based on the standard chemotaxis assay (Figure 1A). We found that consistent with previous findings IAA was strongly attractive and 2-nonanone was strongly repulsive. Strikingly, when these two odorants were placed aside each other, they together repelled the wild-type animals as strongly as 2-nonanone alone (Figures 1B and 1C). To quantify the masking effect of 2-nonanone, we first separately measured the choice index (CI) for 1 μL IAA (CI_Attractant_) or 1 μL 2-nonanone (CI_Repellent_) and the choice index when worms were presented with 1 μL IAA and 1 μL 2-nonanone together (CI_Paring_) as shown in Figure 1A. A positive choice index indicates attraction towards the tested odorant and a negative choice index indicates avoidance. We then calculated the masking index (MI), which was defined as the ratio in percentage of the difference between CI_Pairing_ and CI_Attractant_ to the difference between CI_Repellent_ and CI_Attractant_. Thus, the masking index indicates how much the presence of a repellent, such as 2-nonanone, blocks the attraction of an attractant, such as IAA (Figure 1A and Materials and Methods). Using this paradigm, we found that IAA generated a CI close to 1. In comparison, 2-nonanone generated a CI close to −1 (Figure 1B). Strikingly, paring the two odorants generated a CI close to −1 (Figures 1B and 1C). The results together produced a masking index close to 100%, indicating a near complete masking of IAA by 2-nonanone. To better understand how the indexes were produced by different choices made by the animals in these assays, we also quantified the percentage of the choices made by the worms in each chemotaxis assay (Materials and Methods). We found that more than 92% of the wild-type animals made the right choices in all three behavior assays (Figure 1D). Thus, the presence of 2-nonanone completely masks the attraction of IAA. These results show that under our experimental conditions worms choose to avoid 2-nonanone that signals threat despite the presence of food signals.

**Figure 1.**
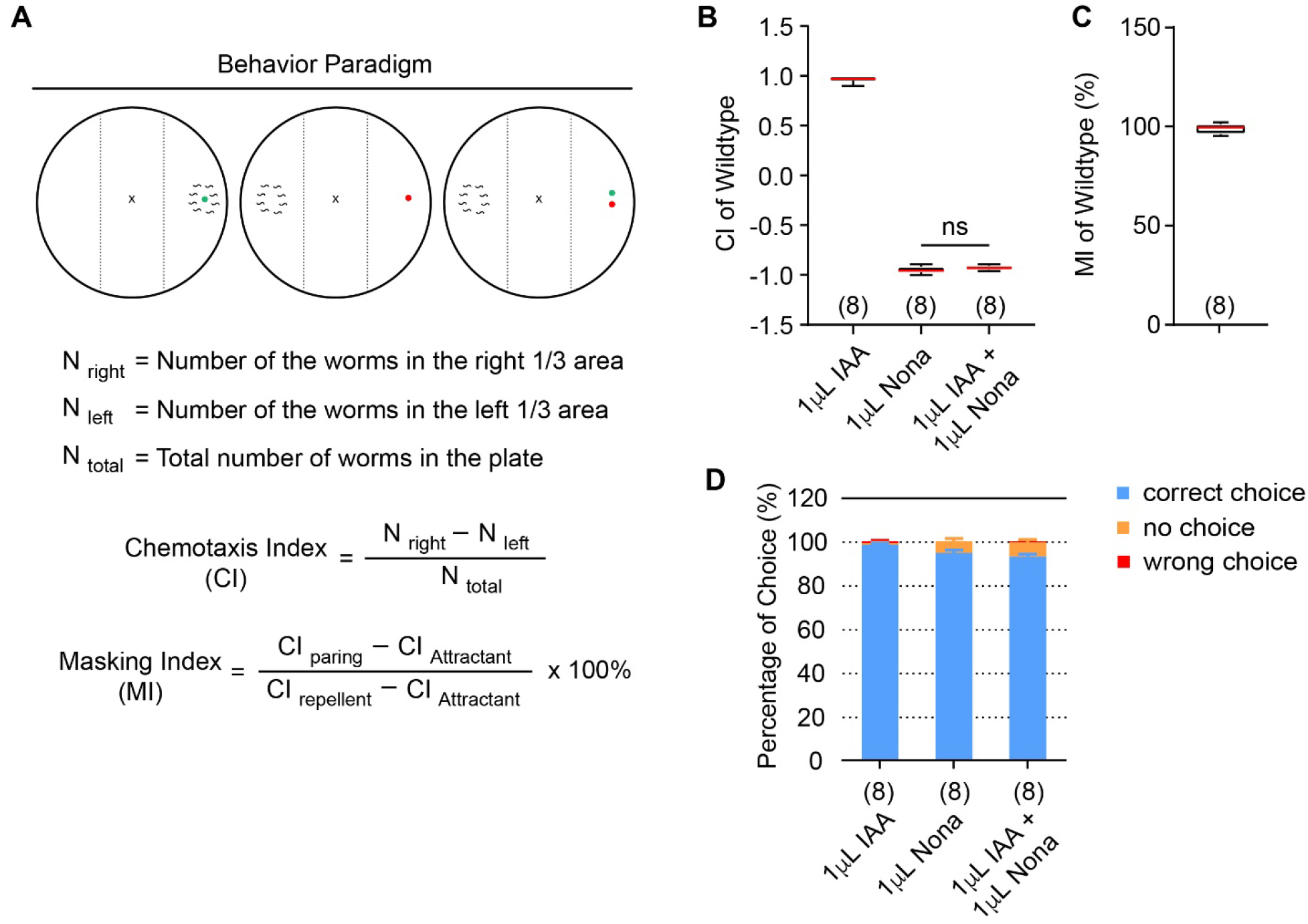
*C. elegans* displays olfactory masking. **(A)** A schematic showing the behavioral assay for chemotaxis assays and the definition of choice index and masking index. **(B, C)** The odorant isoamyl alcohol (IAA) attracts wild-type *C. elegans*, while the odorant 2-nonanone repels it (**B**). However, the presence of 2-nonanone and IAA together repels *C. elegans* similarly as 2-nonanone (**B**), completely masking the attractiveness of IAA (**C**). The numbers of assays are indicated in the parentheses; independent sample *t*-test was used in **B,** because the data is normally distributed; ns, not significant. **(D)** The percentages of choices that generate choice indexes and masking index in **B** and **C.** The numbers of assays are indicated in the parentheses. In **B – C**, box plots indicate median, the first and the third quantile, and the minimal and maximal values; in **D**, mean ± SEM.

### A forward genetic screen identified 3 mutants that are defective in olfactory masking

To understand the regulation of olfactory masking, we conducted a forward genetic screen. We generated a library of mutants by using EMS-based random mutagenesis and screened for F2 clones that were capable of sensing both IAA and 2-nonanone separately, but significantly defective for olfactory masking (Materials and Methods). Using these criteria, we identified 3 mutants, *yx51, yx50* and *yx49* (Figure 2), among ~ 20,000 F2 clones generated by EMS-mutagenesis.

**Figure 2.**
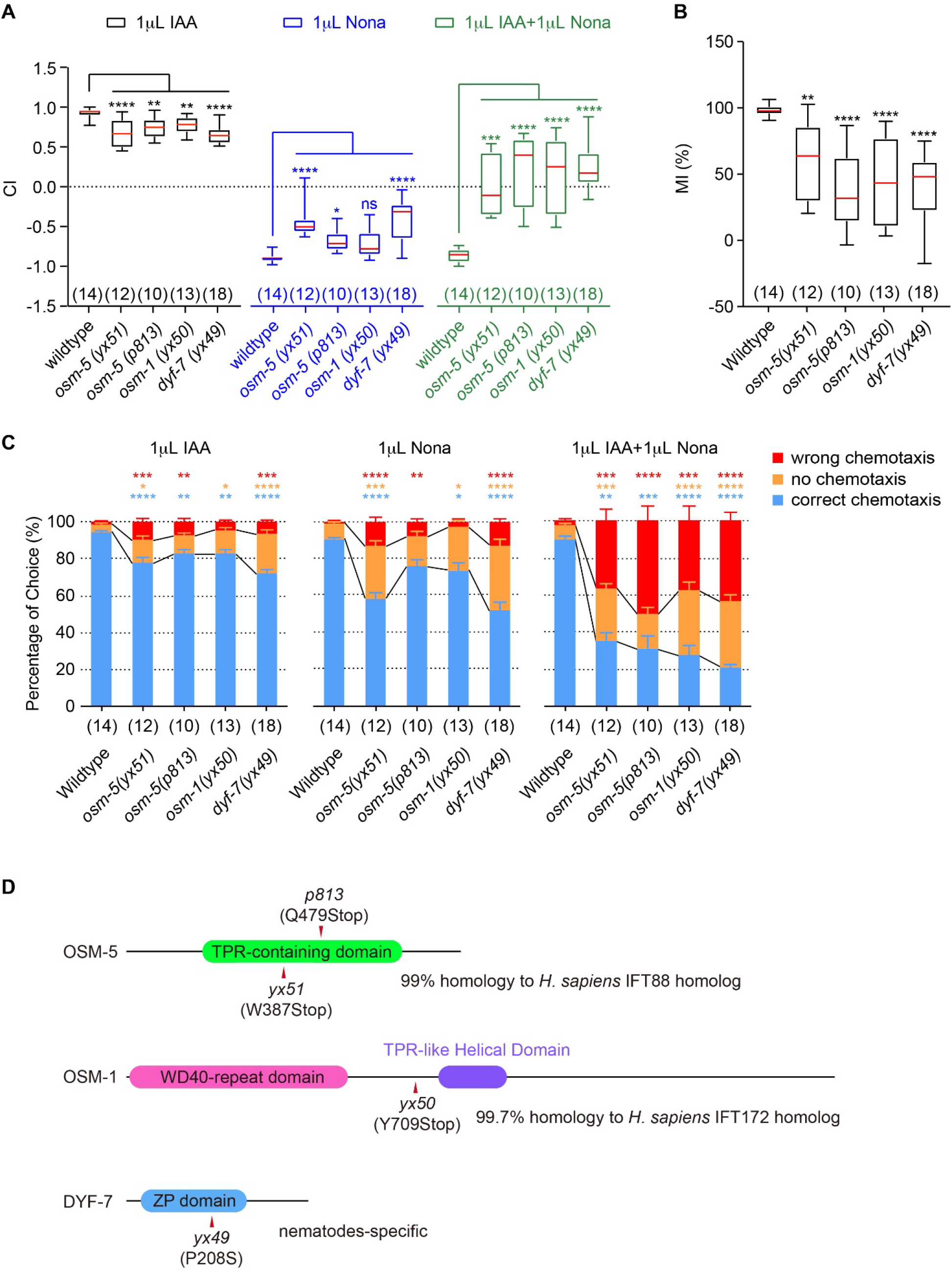
Mutations in genes regulating sensory neurons disrupt olfactory masking. **(A, B)** Mutations in *osm-5, osm-1* and *dyf-7* mildly disrupt chemotaxis to single odorant of IAA or 2-nonanone (**A**), but strongly disrupts olfactory masking (**A, B**). **(C)** The percentages of choices that generate choice indexes and masking indexes in **A, B**. In **A** and **B**, box plots indicate median, the first and the third quantile, and the minimal and maximal values (**A, B**) and in **C**, mean ± SEM. Mutant indexes or percentage were compared with wild type by One way ANOVA with Dunnett multiple comparisons (if the data are normally distributed) or Kruskal-Wallis test with Dunn’s multiple comparisons (if the data are not normally distributed). The numbers of assays are indicated in the parentheses, **** p < 0.0001, *** p < 0.001, ** p < 0.01, * p < 0.05, ns, not significant. **(D)** Schematics showing the proteins domains of OSM-5, OSM-1 and DYF-7, as well as the mutations identified in this or previous studies.

All three mutant animals, *yx51, yx50*, and *yx49*, showed slightly reduced chemotaxis to IAA (Figure 2A, black boxes), and obviously reduced avoidance of 2-nonanone (Figure 2A, blue boxes). The choice indexes for the paring of IAA and 2-nonanone in all three mutants were significantly higher than the choice indexes for 2-nonanone (Figure 2A, green boxes). These results together produced significantly reduced masking indexes for all three mutants (Figure 2B). Because the masking index measures how much paring with 2-nonanone changes the chemotaxis to IAA relative to the difference between the chemotaxis to each of the odorant (Figure 1A), these results indicate that *yx51, yx50* and *yx49* are defective in olfactory chemotaxis and olfactory masking. Although in comparison with wild type, all three mutants showed a significantly decreased portion of animals that made the right choice and an increased portion of the animals that made no choice during chemotaxis to each odorant, they all had a dramatically increased portion of animals that made the wrong choice when tested for olfactory masking (Figure 2C). These results show that significant portions of these mutant animals displayed attraction towards IAA despite the presence of 2-nonanone, which results in defects in generating the masking effect of 2-nonanone on IAA (Figure 2C).

### *osm-5, osm-1*, and *dyf-7* regulate olfactory masking

To identify the genetic lesions that disrupted olfactory masking in *yx51, yx50* and *yx49* mutant animals, we sequenced the genomes of all three mutants, which identified a list of candidate mutations in each mutant strain. We confirmed the causal mutation by testing the rescuing effect of expressing the wild-type candidate genes on olfactory chemotaxis and olfactory masking (Figure 2D). We found that the phenotype of *yx51* was rescued by expressing the wild-type *osm-5* cDNA sequence driven by an *osm-5* promoter (Figures 3A – 3C). Another previously identified mutation in *osm-5, p813*, showed phenotypes similar to that of *yx51. p813* was also rescued by the same DNA construct of *osm-5* (Figures 2A, 2B and 3D – 3F). These results together indicate that the G to A mutation that changes the tryptophan 387 of OSM-5 to a stop codon (Figure 2D) disrupted olfactory masking in the *yx51* mutant animals. Expressing the fosmid WRM0638dE09 that contains the genomic region of *osm-1* rescued the mutant phenotypes of *yx50* (Figures 3G – 3I). WRM0638dE09 also contains another two genes, *pat-9* and *k08b5.2*. However, our whole-genome sequencing analysis did not identify any mutation in the coding region of either of these two genes. Based on all these results, we concluded that the C to A mutation that changes the tyrosine 709 of OSM-1 to a stop codon (Figure 2D) generated the defect in olfactory masking in the *yx50* mutant animals.

**Figure 3.**
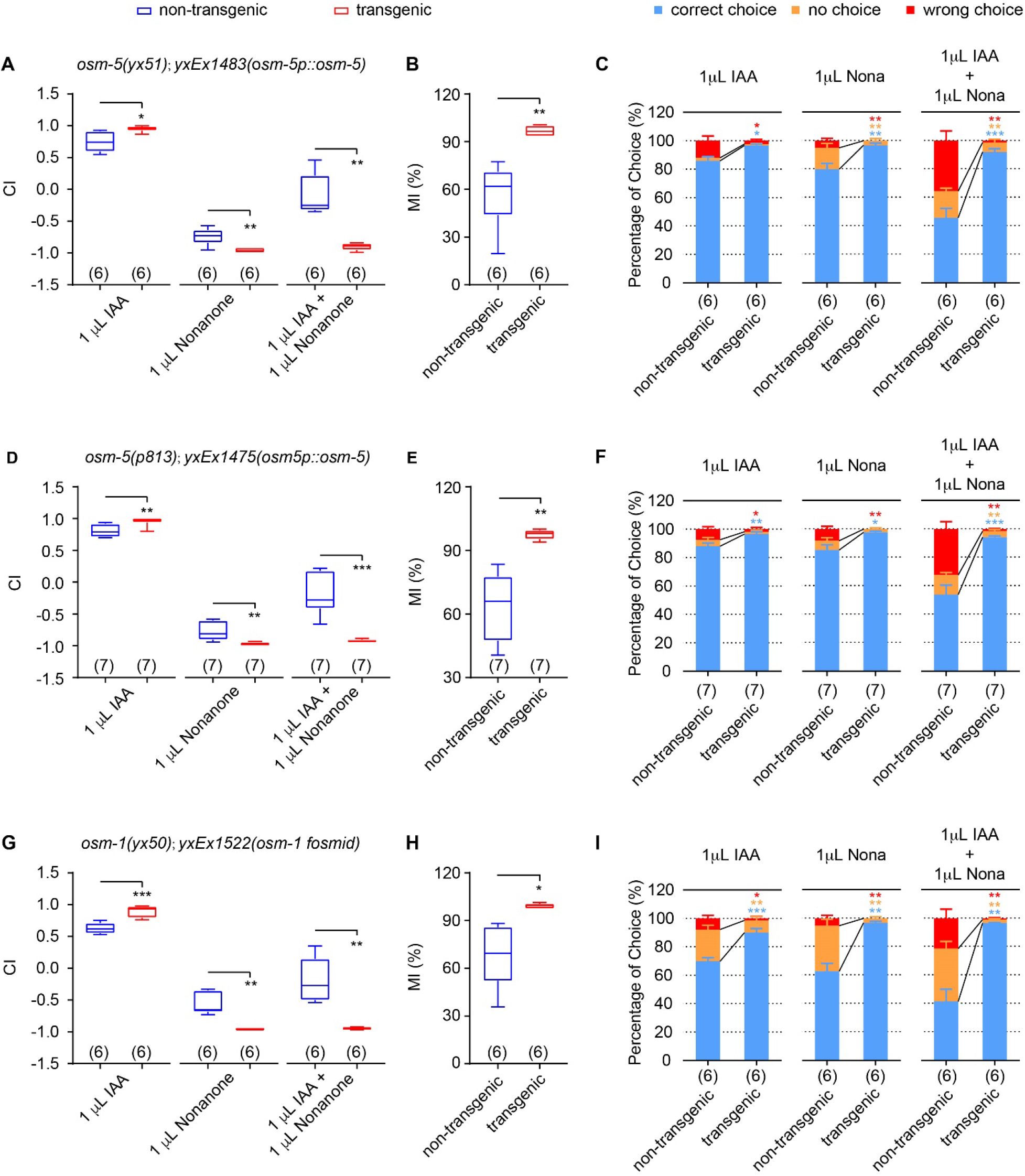
Expressing *osm-5 or osm-1* rescues the defects in olfactory masking. **(A - C)** Expressing a wild-type *osm-5* cDNA with an *osm-5* promoter rescues the defects of *osm-5(yx51)* mutants in chemotaxis to single odorants (**A**) and olfactory masking (**A, B**), as well as the behavioral choices during the assays (**C**). **(D - F)** Expressing a wild-type *osm-5* cDNA with an *osm-5* promoter rescues the defects of *osm-5(p813)* mutants in chemotaxis to single odorants (**D**) and olfactory masking (**D, E**), as well as the behavioral choices during the assays (**F**). **(G - I)** Expressing a wild-type fosmid containing *osm-1* genomic sequence rescues the defects of *osm-1(yx50)* mutants in chemotaxis to single odorants (**G**) and olfactory masking (**G, H**), as well as the behavioral choices during the assays (**I**). In **A, B, D, E, G, H,** box plots indicate median, the first and the third quantile, and the minimal and maximal values, and in **C, F, I,** mean ± SEM. Unpaired t-test (if the data are normally distributed) or Mann-Whitney test (if the data are not normally distributed) was used to compare transgenic animals and their non-transgenic siblings tested in parallel. The numbers of assays are shown in the parentheses, **** p < 0.0001, *** p < 0.001, ** p < 0.01, * p < 0.05, ns, not significant.

Previous studies show that both *osm-5* and *osm-1* encode intraflagellar transport proteins that regulate the generation and function of sensory cilia [25–27]. We speculated that the candidate gene in *yx49* mutant animals may also be related to sensory neurons. Thus, we performed dye-filling assay, which assessed the function of the sensory cilia. We found that *yx49, yx50* and *yx51* were all defective in dye-filling (Supplemental Figure 1), revealing the impaired cilia in these mutant animals. Our sequencing analysis on *yx49* identified a C to T mutation that changed the proline 208 to serine of DYF-7 (Figure 2D). *dyf-7* encodes an extracellular matrix protein that regulates the anchoring of the dendrite tip during the development of sensory neurons [28]. These results suggest that the disfunction of DYF-7 in *yx49* may account for the reduced olfactory masking behavior in this mutant strain. Together, our study identified three genes that are known to regulate cilia structure and function or development of sensory neurons. Similar to previously characterized mutations in these genes [2, 29, 30], *osm-5(yx51), osm-1(yx50)*, and *dyf-7(yx49)* are defective in olfactory chemotaxis (Figure 2A). Our studies characterize the new function of these genes in regulating olfactory masking.

### AWB and ASH neurons function redundantly to regulate olfactory masking

To characterize the mechanism of olfactory masking, we sought the underlying neural circuit. Because two independently generated *osm-5* mutant alleles showed similar phenotypes, we focused on the analysis of *osm-5* hereafter. *osm-5* is expressed in ciliated sensory neurons [25]. Because *osm-5(p813)* animals were defective in DiI staining, which stains several neurons that are exposed to the environment with a red fluorescent lipophilic dye, DiI (Supplemental Figure 1), *osm-5* may function in the DiI labeled amphid chemosensory neurons (ADL, ASH, ASI, ASJ, ASK, AWB) to regulate olfactory masking. We tested the AWB and ASH neurons, because they were shown to respond to 2-nonanone and to mediate 2-nonanone-induced repulsion [4, 20, 21]. We expressed a wild-type *osm-5* cDNA selectively in AWB or ASH in the *osm-5(p813)* mutant animals using cell-specific promoters (Materials and Methods). We found that expressing wild-type *osm-5* in either the AWB neurons using the *str-1* promoter [4] or in the ASH neurons using the *sra-6* promoter [31] significantly rescued the defects of *osm-5(p813)* in olfactory chemotaxis and masking (Figures 4 and Supplemental Figure 2). Together these results identified the critical role of AWB and ASH in regulating olfactory masking.

**Figure 4.**
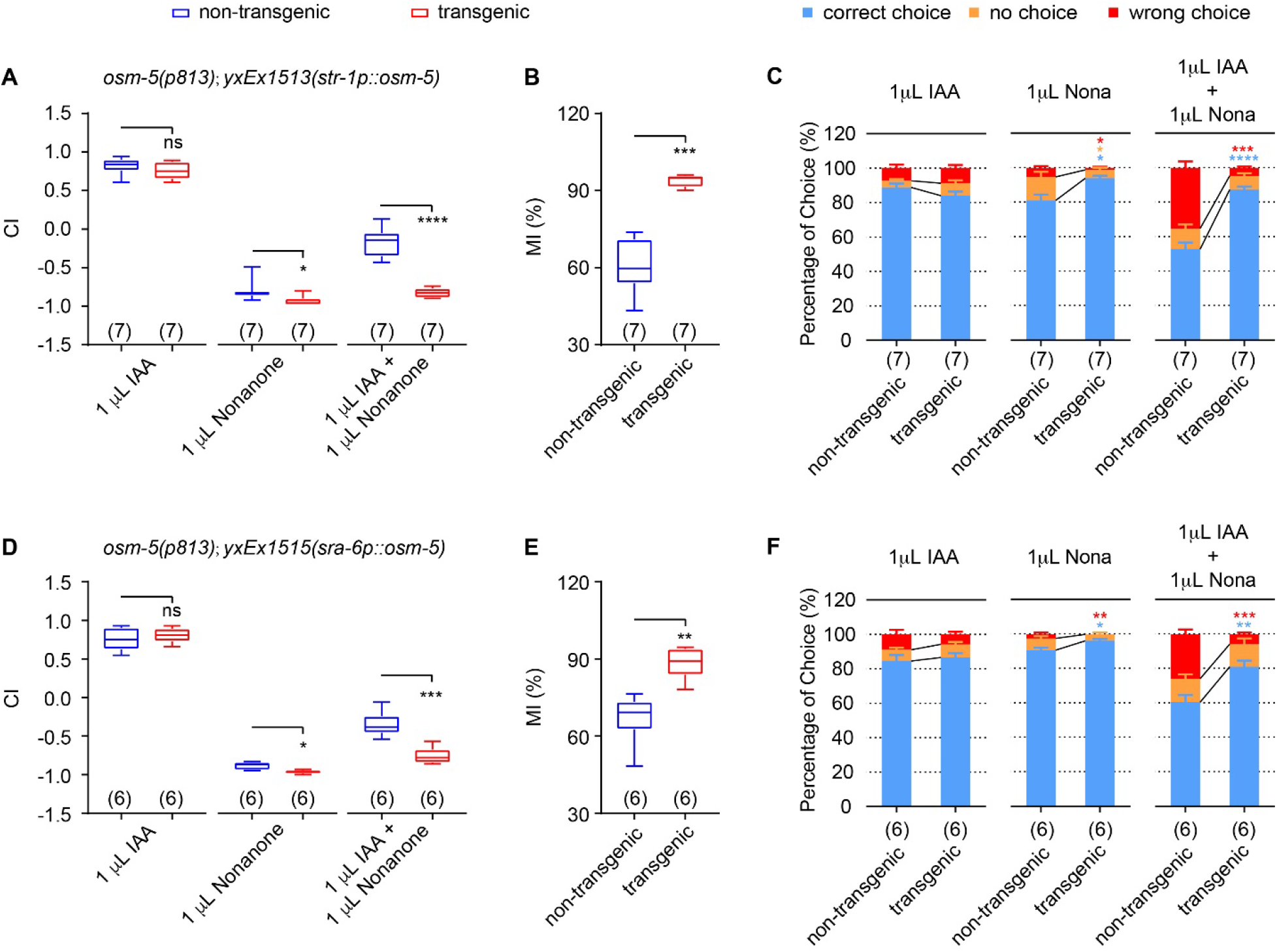
*osm-5* acts in AWB or ASH neurons to regulate chemotaxis and olfactory masking. **(A - C)** Expressing a wild-type *osm-5* cDNA in AWB rescues the defects of *osm-5(p813)* mutants in chemotaxis to single odorants (**A**) and olfactory masking (**A, B**), as well as the behavioral choices during the assays (**C**). **(D - F)** Expressing a wild-type *osm-5* cDNA in ASH rescues the defects of *osm-5(p813)* mutants in chemotaxis to single odorants (**D**) and olfactory masking (**D, E**), as well as the behavioral choices during the assays (**F**). In **A, B, D, E,** box plots indicate median, the first and the third quantile, and the minimal and maximal values, and in **C, F,** mean ± SEM. Unpaired t-test (if the data are normally distributed) or Mann-Whitney test (if the data are not normally distributed) was used to compare transgenic animals and their non-transgenic siblings tested in parallel. The numbers of assays are shown in the parentheses, **** p < 0.0001, *** p < 0.001, ** p < 0.01, * p < 0.05, ns, not significant.

### The AWB neurons function as one of the integration centers for olfactory masking

Next, to understand how the AWB neurons regulate olfactory masking, we measured the response of AWB to IAA or 2-nonanone alone and to the mixture of the two odorants using transgenic animals that selectively expressed a calcium sensor GCaMP6s [32] in AWB. First, we found that exposure to IAA activates AWB, indicated by a rapidly increased GCaMP6 signal that lasted during the presence of IAA stimulation. Removal of IAA led to a gradual decrease in the GCaMP6 signal in AWB (Figures 5A – 5C). Consistent with previous findings, exposure to 2-nonanone modestly suppressed AWB activities and removal of 2-nonanone activated AWB (Figures 5D – 5G). Strikingly, the mixture of IAA and 2-nonanone generated AWB GCaMP6 signal comparable to that generated by 2-nonanone alone (Figures 5H – 5K), showing that at the level of AWB neuronal activities, presence of 2-nonanone masks the effect of IAA stimulation.

**Figure 5.**
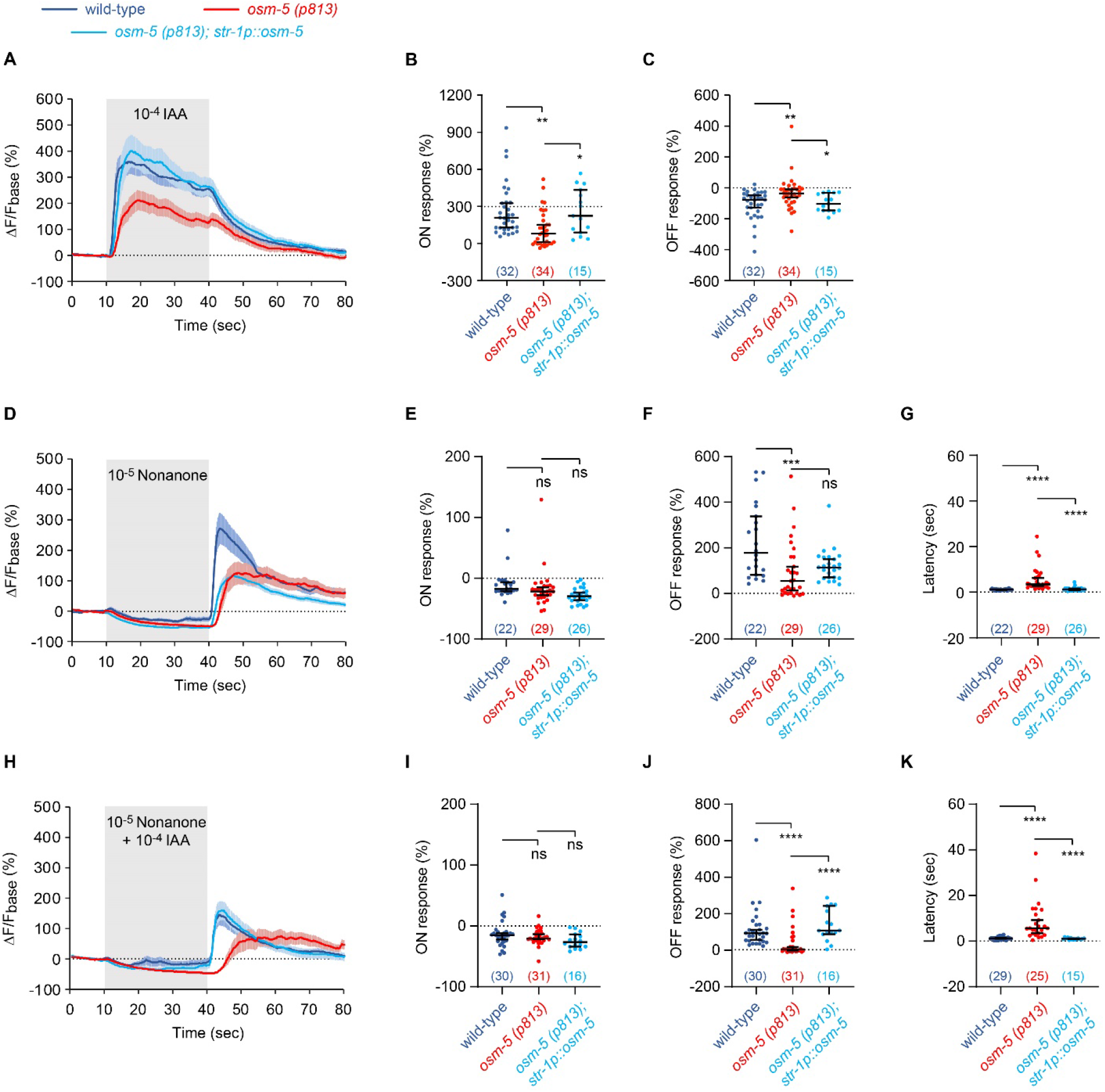
*osm-5* acts in AWB to regulate neuronal responses of AWB to the single odorants and the mixture. **(A - C)** Exposure to IAA increases the GCaMP signal of AWB (ON response) and removal of IAA decreases it (OFF response). The *osm-5(p813)* mutation disrupts both the ON and OFF responses and expressing *osm-5* in AWB rescues the defects. **(D - G)** Exposure to 2-nonanone suppresses the GCaMP signal of AWB (ON response) and removal of 2-nonanone increases it (OFF response). The *osm-5(p813)* mutation decreases the amplitude of OFF response and increases the latency of the OFF response and expressing *osm-5* in AWB rescues the defect in the latency. **(H - K)** Exposure to the mixture of IAA and 2-nonanone suppresses the GCaMP signal of AWB (ON response) and removal of the mixture increases it (OFF response). The *osm-5(p813)* mutation decreases the amplitude of OFF response and increases the latency of the OFF response and expressing *osm-5* in AWB rescues the defects. The change in the fluorescence intensity (ΔF) for each frame is the difference between its fluorescence intensity and the average intensity over the 10-second recording before the stimulus onset (F_base_): ΔF = F - F_base_. The average ΔF/F_base_ % during the 10-second window after onset minus average ΔF/F_base_ % of the 10-second window before onset is used to measure ON response. The average ΔF/F_base_ % during the 10-second window after removal minus average ΔF/F_base_ % of the 10-second window before removal is used to measure OFF response. Latency is defined as the times that it takes for the calcium signal to rise to mean plus 3 × standard deviation. In **A, D, H**, the solid lines and shades are mean and SEM and in **B, C, E – G, I – K**, the horizontal bars in the middle are medians and whiskers cover data points up to 95% confidence interval, individual data points are shown as dots. Kruskal-Wallis Test with Dunn’s multiple comparisons test was used to compare mutants with wild-type and rescued animals, because the data not normally distributed. The numbers of the worms imaged are included in the parentheses. **** p < 0.0001, *** p < 0.001, ** p < 0.01, * p < 0.05, ns, not significant.

Next, we examined the difference between wild type and *osm-5* mutants. First, we found that the amplitude of AWB response to IAA was much lower in *osm-5* than in wild-type, which was rescued by expressing *osm-5* in AWB (Figures 5A – 5C). However, the chemotactic response to IAA was comparable between wild type and *osm-5* (Figure 4A). Therefore, the response of AWB to IAA is unlikely to play a critical role in regulating attraction to IAA, consistent with the critical role of the AWC sensory neurons in mediating the attraction to IAA [23]. Second, we found that the OFF response of AWB to 2-nonanone, *i.e*. the response to the removal of 2-nonanone, displayed a reduced amplitude and increased latency in the *osm-5* mutant animals (Figures 5D – 5G). Expressing *osm-5* in AWB significantly rescued the defect in the latency (Figures 5D and 5G), but not the defect in amplitude (Figures 5D – 5F), which was sufficient to rescue the chemotactic avoidance of 2-nonanone (Figures 4A – 4C and Supplemental Figures 2A – 2C). Thus, the mutation in *osm-5* alters the 2-nonanone avoidance by increasing the latency of 2-nonanone-evoked OFF response. This finding is consistent with previous studies showing that AWB integrates its response to the changes in the odorant concentration over time and triggers escape behavior after the accumulated neuronal response reaches a threshold [21]. Third, we found that when stimulated with a mixture of IAA and 2-nonanone, AWB in *osm-5(p813)* also displayed a decreased amplitude and an increased latency in response to the removal of 2-nonanone, both of which, as well as the olfactory masking effect, were rescued by *osm-5* expression in AWB (Figures 5H – 5K, 4A – 4C and Supplemental Figures 2A – 2C). These results together demonstrate that the latency and amplitude of AWB OFF response regulate olfactory masking.

### ASH neurons respond to 2-nonanone

Previous studies using calcium imaging show that ASH acutely responds to the increase or decrease in the concentration of 2-nonanone [21]. Similarly, we found that exposure to 2-nonanone evoked a rapid increase in the GCaMP6 signal of ASH and removal of 2-nonanone triggered a rapid decrease (Figures 6A – 6C). The mutation in *osm-5(p813)* disrupted the responses of ASH to the onset and removal of 2-nonanone. Both of the defects in ASH neuronal responses, as well as the behavioral defects of the *osm-5(p813)* mutants in their chemotaxis to 2-nonanone and olfactory masking, were rescued by expressing the wild-type *osm-5* in ASH (Figures 6A – 6C, 4D – 4F and Supplemental Figures 2D – 2F). Together, these results indicate that ASH regulates olfactory masking by responding to 2-nonanone.

**Figure 6.**
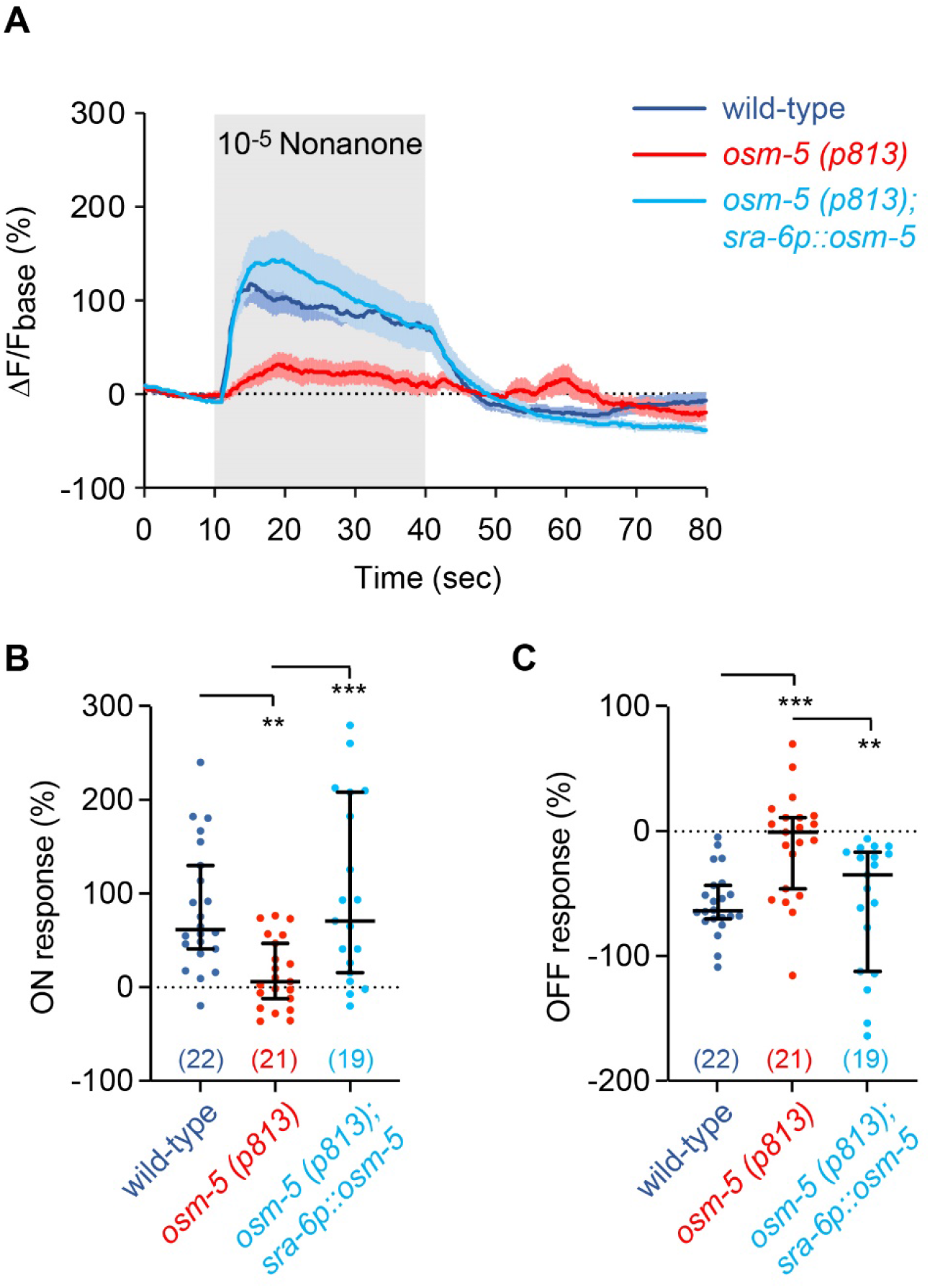
*osm-5* acts in ASH to regulate neuronal response of ASH to 2-nonanone. **(A - C)** Exposure to 2-nonanone increases the GCaMP signal of ASH (ON response) and removal of 2-nonanone decreases it (OFF response). The *osm-5(p813)* mutation disrupts both the ON and OFF responses and expressing *osm-5* in ASH rescues the defects. The change in the fluorescence intensity (ΔF) for each frame is the difference between its fluorescence intensity and the average intensity over the 10-second recording before the stimulus onset (F_base_): ΔF = F - F_base_. The average ΔF/F_base_ % during the 10-second window after onset minus average ΔF/F_base_ % of the 10-second window before onset is used to measure ON response. The average ΔF/F_base_ % during the 10-second window after removal minus average ΔF/F_base_ % of the 10-second window before removal is used to measure OFF response. In **A,** the solid lines and shades are mean and SEM and in **B, C,** the horizontal bars in the middle are medians and whiskers cover data points up to 95% confidence interval, individual data points are shown as dots. One way ANOVA with Dunnett multiple comparisons (if the data are normally distributed) or Kruskal-Wallis test with Dunn’s multiple comparisons (if the data are not normally distributed) was used to compare mutants with wild-type and rescued animals. The numbers of the worms imaged are included in the parentheses. *** p < 0.001, ** p < 0.01.

### The *osm-5(p813)* mutant animals are defective in several complex olfactory tasks

Finally, we tested how general the *osm-5(p813)* mutant animals were defective in olfactory masking by examining the masking effect of 2-nonanone on another commonly used attractant, benzaldehyde [2]. We paired 0.02 μL benzaldehyde with 1 μL 2-nonanone. The wild-type chemotaxis to simultaneous presence of benzaldehyde and 2-nonanone is indistinguishable from that to 2-nonanone alone (Figure 7A), demonstrating a complete masking of the attraction to benzaldehyde by 2-nonanone. While the *osm-5(p813)* mutant animals showed normal chemotactic behavior towards benzaldehyde and slightly reduced chemotaxis towards 2-nonanone, they are dramatically defective in response to the paired odorants (Figure 7A). These results indicate that *osm-5(p813)* mutant animals are also defective in olfactory masking of benzaldehyde by 2-nonanone.

**Figure 7.**
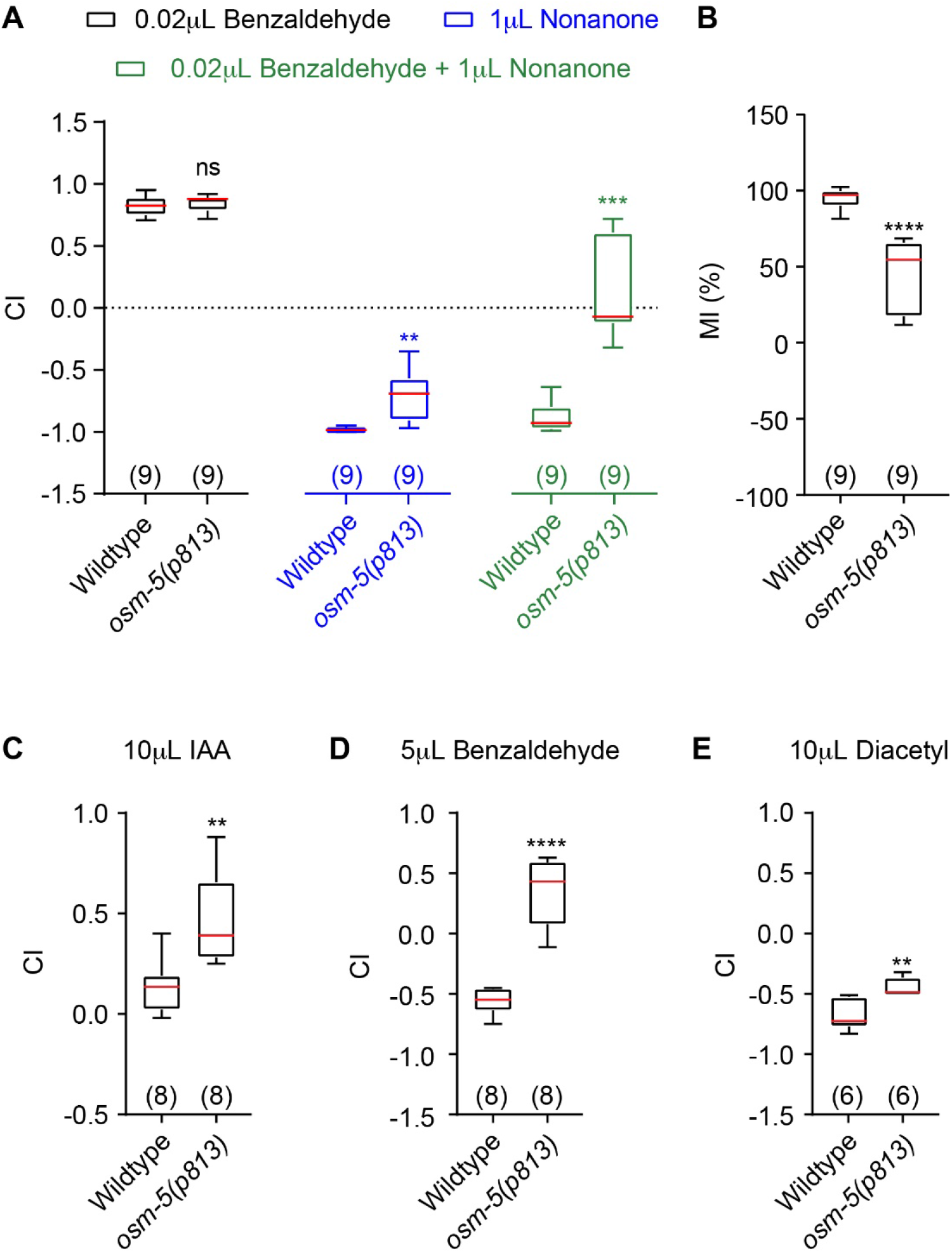
The *osm-5(p813)* mutant animals are defective in several olfactory tasks. **(A, B)** The *osm-5(p813)* mutant animals are defective in olfactory masking of benzaldehyde by 2-nonanone. Sample *t* test (if the data are normally distributed) or Mann-Whitney test (if the data are not normally distributed) was used to compare the mutant indexes with the wild-type indexes. **(C - E)** The *osm-5(p813)* mutant animals are defective in high concentration-dependent repulsion of IAA, benzaldehyde and diacetyl. Unpaired *t* test is used to compare mutant choice indexes with wild-type choice indexes, because the data are normally distributed. For all, box plots indicate median, the first and the third quantile, and the minimal and maximal values. The numbers of assays are shown in the parentheses, **** p < 0.0001, *** p < 0.001, ** p < 0.01, * p < 0.05, ns, not significant.

Previously, it was shown that increasing the concentration of IAA switches the odorant from attractive to repulsive [33]. At lower concentrations, exposure to IAA suppresses AWC calcium activities and removal of IAA activates AWC [34]. At higher concentrations, exposure to IAA activates the nociceptive sensory neuron ASH and removal of IAA activates AWB similarly as the repellent 2-nonanone [4, 33]. Because the mutation in *osm-5(p813)* disrupted the neuronal response of AWB and ASH to repulsive odorants and the mixture of attractive and repulsive odorants, we hypothesized that *osm-5(p813)* also disrupted the repulsion of IAA at higher concentrations. We found that in comparison with wild type, *osm-5(p813)* mutant animals are more attracted to not only 10μL IAA, but also to 5μL benzaldehyde or 10μL diacetyl (Figures 7C – 7E). These results indicate that *osm-5(p813)* also disrupts concentration-dependent chemotaxis towards odorants. Together, these results suggested that OSM-5 regulates several complex olfactory tasks.

## DISCUSSION

In this study, we developed a new behavior paradigm to study olfactory masking using a binary mixture of two odorants of opposing valences, the repulsive odorant 2-nonoanone and the attractive odorant IAA that respectively represent danger and reward. Wild-type animals show a near complete masking effect of 2-nonanone on IAA. Mutations in three genes, *osm-5, osm-1, dyf-7* that are known to regulate the development and function of sensory cilia, show an impaired masking effect when 2-nonanone is paired with different attractants. By analyzing the function of *osm-5*, we found that OSM-5 acts in either AWB or ASH, two pairs of sensory neurons that detect repellents and mediate avoidance response, to regulate olfactory masking. AWB integrate the sensory responses to 2-nonanone and IAA to generate the cellular basis for olfactory masking.

### The sensory cilia are important for the olfactory masking effect

The cilia of the *C. elegans* sensory neurons play an important role in sensing external cues. As major sensory machineries, the cilia contain sensory receptors, signaling proteins and display specialized morphologies [35]. Thus, an intact cilium is important for the sensorimotor behaviors of the worms, including olfaction. Previous studies have shown that mutation disrupting cilia structure, such as those in *bbs-7* or *bbs-8* or *odr-3*, reduce chemotaxis to odorants, including IAA [36, 37]. By analyzing about 20,000 mutants generated from random mutagenesis, we identified 3 mutations that disrupted the olfactory masking effect of 2-nonanone on attractants and all 2 mutations were mapped in the genes involved in ciliogenesis. These results reveal that the normal function of the sensory cilia is important for generating an olfactory masking, besides its known function in olfactory sensation.

### Redundant circuits are used for olfactory masking

Instinctive behaviors, such as feeding and avoiding dangers, are essential for survival of the animals. Thus, the underlying neural circuits need to be able to generate robust behavioral outputs. This demand is partly met through the redundancy of the circuits. For example, subsets of the Agouti related protein-expressing neurons in the brain form redundant circuits, each of which is sufficient to regulate the feeding behavior in mice [38]. However, anatomically redundant neurons do not necessarily lead to functional redundancy. For example, the sister mitral cells innervating the same glomerulus in the mammalian olfactory bulb generate different spike patterns in response to the same sensory input [39]. Thus, the functional redundancy cannot be predicted based on anatomical properties. Previous studies show that both AWB and ASH sensory neurons regulate the avoidance of 2-nonanone, but in different ways. AWB responds to a decrease in 2-nonanone concentration with increased intracellular calcium transients, integrates the signal over time, and suppresses reversals when the integrated signal reaches a threshold. In contrast, ASH responds to an increase in 2-nonanone concentration to trigger reversals upon the response [21]. Thus, it is likely that AWB and ASH neurons signal to separate downstream circuits to process information of 2-nonanone and generate behavioral outputs. In our study, we observed that restoring the expression of *osm-5* in either AWB or ASH neurons almost fully rescued the olfactory masking effect of 2-nonanone, suggesting their redundant function in this olfactory task. Because *C. elegans* showed instinctive avoidance to 2-nonanone, this circuit redundancy may be important to generate reliable avoidance of dangers in presence of food signals in order to benefit survival. The redundancy is also reflected in how they rescued the behavioral outcome in individual worms. We divided the worms in each chemotaxis plate to 3 groups, the group of correct choice, no choice, and wrong choice, and found that either AWB or ASH expression of *osm-5* reduced the percentage of the worms that made wrong choice, but without significantly reducing the percentage of the worms that made no choice. Because expressing *osm-5* with its own promoter significantly reduced the percentage of the worms that made a wrong choice as well as worms that made no choice in the *osm-5* mutants, we speculate that AWB and ASH neurons work together to fully rescue the behavior, or additional *osm-5*-expressing sensory neurons also contribute to the regulation of olfactory masking.

### Olfactory masking is a context-dependent response to odorants

While the molecular and cellular mechanisms underlying olfactory sensation have been well studied using individual chemicals, the olfactory stimuli that an animal encounters in its daily life are often mixtures of different odorant chemicals. Previous studies in *Drosophila* show that mixing an aversive odorant with an attractive odorant can significantly reduce the behavioral attraction to the attractive odorant. The context-dependent response to the attractant is regulated by lateral inhibition to the glomerulus that responds to the attractant [40]. In mice, different ligands for the same trace amine-associated receptor can elicit different behavior, likely due to different additional receptors activated by each of these ligands [3]. These context-dependent effects on behavioral response to the odorants provide mechanisms through which the nervous system processes and integrates complex olfactory information. Here, we show that presence of the repulsive odorant 2-nonanone completely masks the attraction of IAA in behavior and the activation of AWB neurons by IAA. These results show that when 2-nonanone, which indicates danger, is present as a context, both the behavior and neuronal response to IAA are modulated. It is conceivable that these context-dependent regulations of olfactory response were built into the nervous system of the worm to ensure the avoidance of danger-associated signals.

## Materials and Methods

### Strains and transgenes

*C. elegans* strains were maintained under standard conditions at 20°C [41]. Hermaphrodites were used in the study. The strains that were used include: N2, ZC2925 *dyf-7(yx49)* X, SP1196 *dyf-7(m539)* X, ZC2926 *osm-1(yx50)* X, PR808 *osm-1(p808)* X, PR816 *osm-1(p816)* X, ZC2927 *osm-5(yx51)* X, PR813 *osm-5(p813)* X, ZC2858 *osm-5(p813)* X; *yxEx1475[osm-5p::osm-5; unc-122p::GFP]*, ZC2868 *osm-5(yx51)* X; *yxEx1483[osm-5p::osm-5; unc-122p::GFP]*, ZC2880 *osm-5(p813)* X; *yxEx1489[str-1p::osm-5; unc-122p::GFP]*, ZC2910 *osm-5(p813)* X; *yxEx1513[str-1p::osm-5; unc-122p::GFP]*, ZC2911 *osm-5(p813)* X; *yxEx1514[sra-6p::osm-5; unc-122p::GFP]*, ZC2912 *osm-5(p813)* X; *yxEx1515[sra-6p::osm-5; unc-122p::GFP]*, ZC2919 *osm-1(yx50)* X; *yxEx1522[Fosmid WRM0638dE09; unc-122p::GFP]*, ZC2904 *yxEx1507[str-1p::GCaMP6s; unc-122p::DsRed]*, ZC2921 *osm-5(p813)* X; *yxEx1507[str-1p::GCaMP6s; unc-122p::DsRed]*, ZC2922 *unc-13(e51)* I; *yxEx1507[str-1p::GCaMP6s; unc-122p::DsRed]*, ZC2905 *yxEx1508[sra-6p::GCaMP6s; unc-122p::DsRed]*, ZC2923 *osm-5(p813)* X; *yxEx1508[sra-6p::GCaMP6s; unc-122p::DsRed]*, ZC2924 *unc-13(e51)* I; *yxEx1508[sra-6p::GCaMP6s; unc-122p::DsRed]*, ZC3170 *osm-5(p813)* X; *yxEx1643[str-1p::GCaMP6s; str-1p::osm-5; unc-122p::GFP]*, ZC3148 *osm-5(p813)* X; *yxEx1508[sra-6p::GCaMP6s; unc-122p::DsRed]; yxEx1514[sra-6p::osm-5; unc-122p::GFP]*.

To generate transgenic animals, fosmid WRM0638dE09 (Wellcome Trust Sanger Institute) was injected into *yx50* at 5 ng/μL. The 242 bp 5’ upstream sequence of *osm-5* was generated by PCR (Primers: 5’-tttattgttttgaaattgaaagactcg-3’ and 5’-taagaaaagtgttctcagaagaaatagag-3’) and cloned into PCR8 to generate the entry clone (pCR8-osm-5p). The *osm-5* cDNA sequence and GCaMP6s coding sequence were used to generate the destination vector pDEST-*osm-5* and pDEST-GCaMP6s by Gateway vector conversion kit. To generate *osm-5p::osm-5*, Gateway LR reaction was performed between pDEST-osm-5 and PCR8-osm-5p. *osm-5p::osm-5* was injected into *yx51* or *p813* at 36 ng/μL. The 4700 bp 5’ upstream sequence of str-1 [4] was cloned into pCR8 to generate the entry clone (PCR8-str-1p). The 3269 bp 5’ upstream sequence of sra-6 [31] was cloned into pCR8 to generate the entry clone (PCR8-sra-6p). To generate plasmids *str-1p::osm-5* (37 ng/μL or 5 ng/μL), *sra-6p::osm-5* (5 ng/μL), *str-1p::GCaMP6s* (25 ng/μL), *sra-6p::GCaMP6s* (34 ng/μL), Gateway LR reactions were performed between above mentioned corresponding destination and entry vectors. Microinjection (at 100 ng/μL total DNA concentration with PUC19 added when needed) was performed as described previously [42] with either *unc-122p::GFP* or *unc-122p::DsRed* at 30 ng/μL as a co-injection marker.

### Behavior assay

One-day old adult animals (more than 50 worms for each assay) were washed 4 times by M9 buffer (3g/L KH_2_PO_4_, 6g/L Na_2_HPO_4_, 5g/L NaCl, 0.12g/L MgSO_4_), and then placed at the center (marked with a “X” in Figure1A) of a 10 cm NGM-agar (nematode growth medium) plate. The odorants were placed 3.33 cm away from the center of the 10 cm NGM-agar plate and the control (NGM buffer) was on the opposite side. 1 μL of 1 mol/L sodium azide was placed beside the odorants and on the other side of the plate equidistant from the center to immobilize the worms that reach the odorant or the control. The extra M9 buffer was then removed by Kimwipes. The plate was sealed by Parafilm and placed upside-down on the bench for 1 hour. The numbers of worms in all three areas, as shown in Figure 1A, were counted manually, and the chemotaxis index (CI) and the masking index (MI) were calculated according to the formulas in Figure 1A. The worms stayed in the middle area were the ones made no choice. The worms moved to the area with odor(s) were the ones made correct choice to IAA, and wrong choice to 2-nonanone or IAA + 2-nonanone. The worms stayed in the area furthest away from the odor(s) were the ones made wrong choice to IAA, and correct choice to 2-nonanone or IAA + 2-nonanone. Statistical analyses were performed using GraphPad Prism.

### Mutagenesis, screen, and mutant identification

To generate mutants for forward genetic screen, L4-stage wild-type hermaphrodites (P0) were treated with 0.5% ethanemethylsulfonate (EMS diluted with M9 buffer) for 4 hours and washed with M9 buffer for four times. After recovering on a regular cultivation plate for overnight, 200 EMS-treated P0 worms were transferred to 50 fresh cultivation plates (4 worms/plate) to reproduce and removed after around 50 eggs (F1s) were found on each plate. F1 worms were removed after 200 ~ 500 eggs (F2) were found on each plate and F2 worms were tested for olfactory masking. The F2 worms stayed on the side of the odorant mixture were collected and cultivated into individual F2 clones for future analysis. Mutants that were morphologically defective, or severely uncoordinated in locomotion, or retarded in development, or strongly defective in single odorant chemotaxis were excluded. Based on these criteria, 3 F2 clones, *yx49, yx50* and *yx51*, were identified as mutants for olfactory masking.

All three mutants were outcrossed three to four times with a wild-type genetic background and sequenced by Illumina Hi-Seq 2500 (single-read, 50-basepair read length). Sequencing reads were aligned to the WS260 reference genome and analyzed by MiModD (http://mimodd.readthedocs.io/en/latest/). The function of candidate genes, suggested by MiModD, in olfactory masking was tested by using the existing mutant allele(s) for the genes to see if they phenocopied the mutants identified from EMS-mutagenesis. If no mutant was available for a gene of interest, a fosmid containing the gene of interest was tested for its rescuing effect.

### Calcium imaging

Calcium imaging was performed using microfluidic device essentially as we previously published [43–45]. Briefly, fresh solutions were prepared before each recording session by dissolving the tested chemical(s) in nematode growth medium buffer (3 g/L NaCl, 1 mM CaCl_2_, 1 mM MgSO_4_, 25 mM potassium phosphate buffer pH6.0) to the indicated concentration(s). Fluorescence time-lapse imaging was recorded using a Nikon Eclipse Ti-U inverted microscope with a 40X oil immersion objective and a Yokogawa CSU-X1 scanner unit and a Photometrics CoolSnap EZ camera at 5 frames per second. The GCaMP signal from the soma of AWB or ASH was measured using Fiji. The change in the fluorescence intensity (ΔF) for each frame was the difference between its fluorescence intensity and the average intensity over the 10-second recording before the stimulus onset (F_base_): ΔF = F - F_base_. To analyze the response evoked by the onset of the stimulus for each genotype (ON response), the average ΔF/F_base_ % during the 10-second window after onset minus average ΔF/F_base_ % of the 10-second window before onset was quantified. To analyze the response evoked by the removal of the stimulus (OFF response), the average ΔF/F_base_ % of the 10-second window after removal minus the average ΔF/F_base_ % of the 10-second window before removal was quantified. The latency for the OFF response was defined as the time needed for the calcium signal to reach mean + 3 x SD (standard deviation). The statistical methods are shown in the figure legends.

## ACKNOWLEDGEMENTS

Some strains were provided by the CGC, which is funded by NIH office of Research Infrastructure Program (P40 OD10440). We thank Dr. Sengupta P. for sharing a plasmid containing *osm-5* cDNA sequence.

## DATA REPORTING

All data supporting the manuscript are included in the figure and supporting figures. The genome sequencing results are deposited at: https://dataview.ncbi.nlm.nih.gov/object/PRJNA720651?reviewer=ghdejudsbkeo3v4vioukiq0u91

## Supporting information for

**Figure S1.**
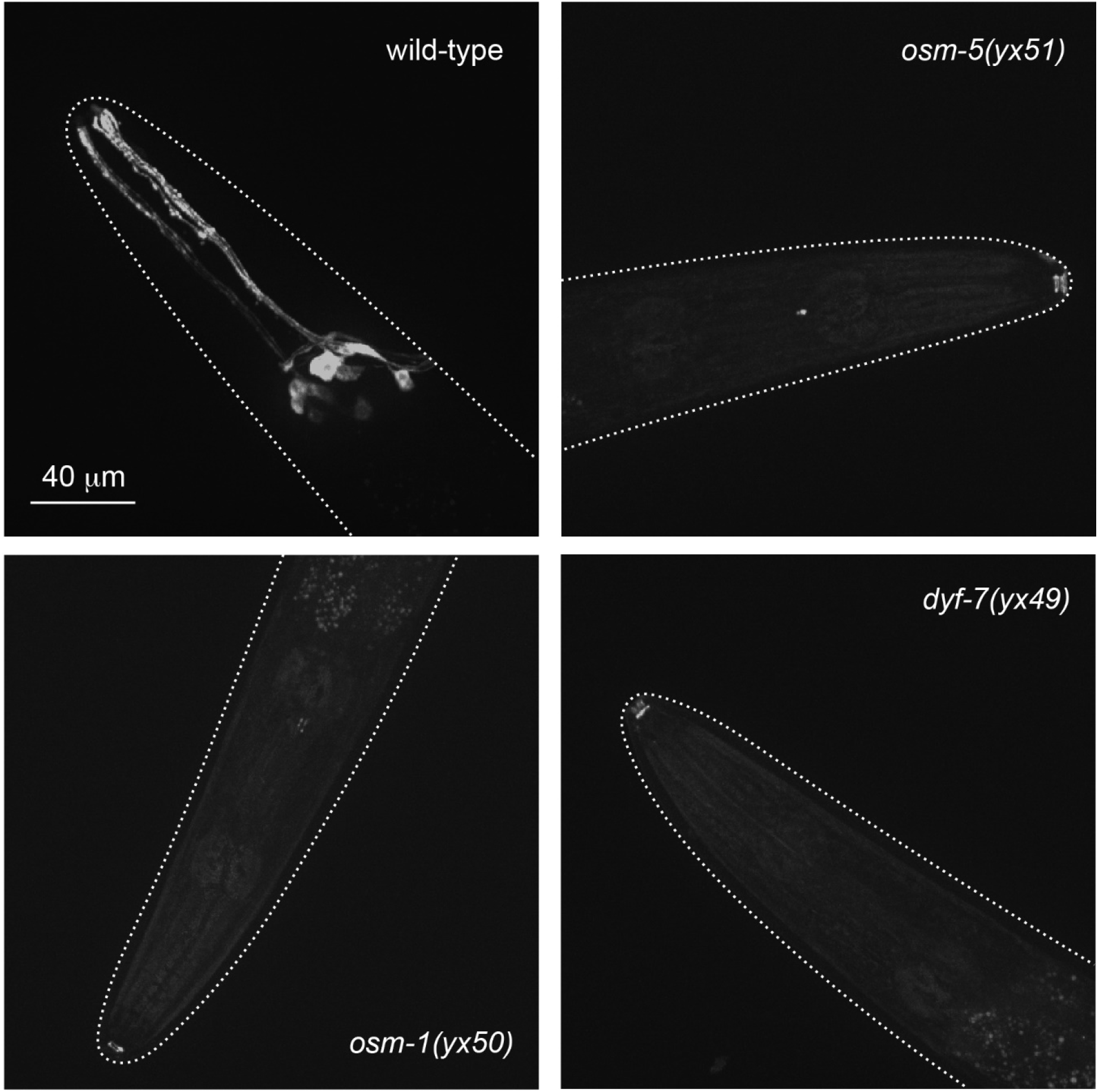
Mutants defective in olfactory masking are defective in DiI staining. Representative images of wild type and olfactory masking mutants after dye fill of DiI. Several neurons in wild type uptake DiI from the environment and generate fluorescent signals. In contrast, the *yx51, yx50* and *yx49* mutants do not show a dye fill signal.

**Figure S2.**
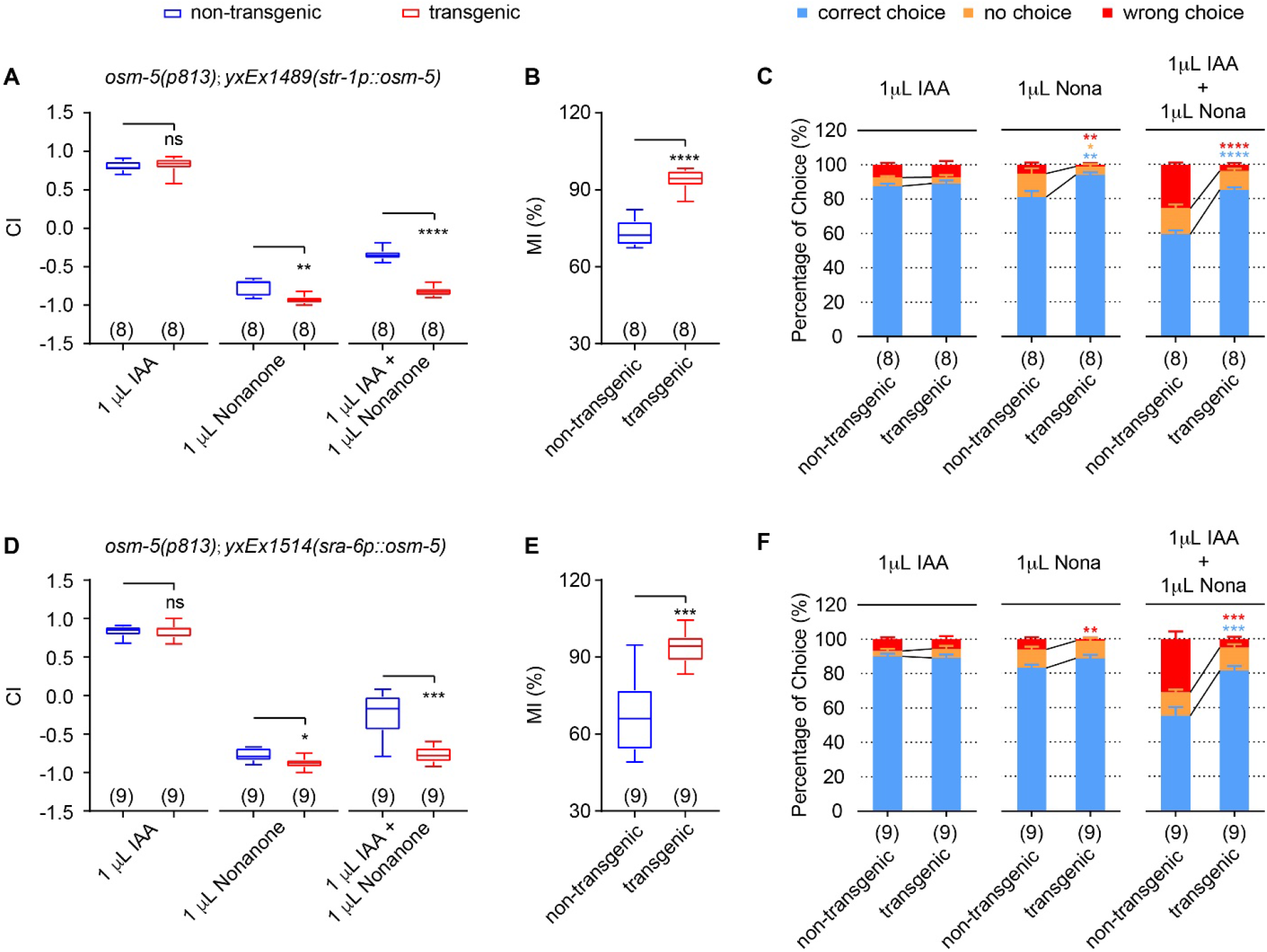
The results from additional transgenic lines generated independently from those in Figure 4 show that *osm-5* acts in the AWB or ASH neurons to regulate chemotaxis and olfactory masking. **(A - C)** Expressing a wild-type *osm-5* cDNA in AWB rescues the defects of *osm-5(p813)* mutants in chemotaxis to single odorants (**A**) and olfactory masking (**A, B**), as well as the behavioral choices during the assays (**C**). **(D - F)** Expressing a wild-type *osm-5* cDNA in ASH rescues the defects of *osm-5(p813)* mutants in chemotaxis to single odorants (**D**) and olfactory masking (**D, E**), as well as the behavioral choices during the assays (**F**). In **A, B, D, E,** box plots indicate median, the first and the third quantile, and the minimal and maximal values, and in **C, F,** mean ± SEM. Unpaired t-test (if the data are normally distributed) or Mann-Whitney test (if the data are not normally distributed) was used to compare transgenic animals and their non-transgenic siblings tested in parallel. The numbers of assays are shown in the parentheses, **** p < 0.0001, *** p < 0.001, ** p < 0.01, * p < 0.05, ns, not significant.

